# Wide-Field Calcium and Flavoprotein Autofluorescence Imaging in Living Mice

**DOI:** 10.64898/2026.05.14.725112

**Authors:** Takamasa Yoshida

## Abstract

Wide-field imaging (WFI) is a mesoscopic approach for monitoring cortex-wide activity with high temporal resolution and a broad field of view. Owing to its simple optical configuration and compatibility with chronic preparations, WFI has become an important tool in systems neuroscience and disease-model research. In this chapter, we describe practical protocols for chronic transcranial WFI in mice using two complementary optical signals: genetically encoded calcium indicators (GCaMP) and endogenous flavoprotein autofluorescence. Calcium imaging provides a robust readout of neuronal population activity, whereas flavoprotein imaging reflects mitochondrial redox dynamics and cellular metabolic demand. We detail procedures for animal preparation, skull clearing, headplate implantation, macroscope assembly, synchronized sensory stimulation, triggered image acquisition, and MATLAB-based data analysis. The analysis workflow includes ΔF/F normalization, reference-based signal correction, and artifact reduction, followed by trial averaging, atlas registration, and region-of-interest analysis. Because imaging is performed through the intact skull, the protocol enables repeated longitudinal measurements in the same animal over extended periods. This approach is reproducible, cost-effective, and adaptable to studies of cortical physiology and neurological disorders.

## 1. Introduction

Wide-field imaging (WFI) is a mesoscopic approach for monitoring cortex-wide neural activity in vivo with high temporal resolution and a broad field of view [1–3]. Unlike point-scanning microscopy, WFI enables simultaneous recording from multiple cortical regions across one or both hemispheres, thereby allowing the investigation of distributed sensory, motor, and cognitive processes in behaving animals. Its relatively simple optical configuration also renders WFI an accessible and cost-effective platform for systems neuroscience [4,5].

Genetically encoded calcium indicators (GECIs), such as GCaMP, serve as a robust optical proxy for neuronal population activity by converting intracellular calcium transients into fluorescence signals [6,7]. Combined with transgenic strategies or viral expression methods, GECI-based WFI supports stable cortex-wide recordings in awake mice and has been widely used to investigate large-scale dynamics during learning, decision-making, movement, spontaneous behavior, and multisensory integration [8–14].

A complementary signal source is endogenous flavoprotein autofluorescence, which reflects mitochondrial redox dynamics and oxidative metabolism, offering an intrinsic readout of cellular energy demand [15]. Because it requires neither genetic modification nor exogenous dyes, flavoprotein imaging is well suited for wild-type animals and disease models. Intact-skull transcranial imaging has been employed for mesoscale mapping of sensory responses and cortical plasticity [16,17], and recent protocols have systematized practical implementation [18].

Together, calcium and flavoprotein imaging provide complementary information on neuronal activity and neurometabolic function. Their combined use can help identify circuit-level dysfunction in neurological disorders. Chronic WFI also allows repeated longitudinal measurements in the same animal and has been applied to models of stroke, Alzheimer’s disease, autism spectrum disorder, experimental parkinsonism, stress-related conditions, and other brain disorders [19–34]. Such longitudinal cortex-wide imaging may be particularly valuable for Parkinson’s disease (PD), in which abnormalities extend beyond nigrostriatal degeneration to include cortical network dysfunction and mitochondrial impairment [35,36].

In this chapter, we describe a practical protocol for chronic transcranial WFI in mice using two complementary approaches: (1) GCaMP-based calcium imaging in transgenic mice and (2) flavoprotein autofluorescence imaging in wild-type mice. We outline procedures for animal preparation, optical hardware, sensory stimulation, image acquisition, and MATLAB-based analysis. The protocol is designed to be reproducible, affordable, and adaptable to studies of cortical physiology, neurological disorders, and experimental models relevant to PD.

## 2. Materials

### 2.1 Animals

1. Wild-type C57BL/6J mice for flavoprotein autofluorescence imaging experiments.
2. Thy1-GCaMP6s transgenic mice (GP4.3 line; RRID:IMSR_JAX:024275) for calcium imaging experiments. This line expresses GCaMP6s in neuronal populations under the Thy1 promoter [37].

All animal procedures were approved by the Teikyo University Animal Ethics Committee (No. 22-018) and were conducted in accordance with institutional guidelines and national regulations for laboratory animal welfare.

### 2.2 Reagents for Surgery and Animal Preparation

1. Mixed anesthetic solution: medetomidine hydrochloride, midazolam, and butorphanol tartrate diluted in sterile saline.
2. Atipamezole hydrochloride solution: anesthetic reversal agent.
3. Lidocaine jelly, 2%: local analgesic.
4. Sterile normal saline.
5. Ophthalmic ointment: corneal protection.
6. Depilatory cream.
7. Chlorhexidine disinfectant solution.
8. 80% ethanol.
9. Alginate wound dressing material.
10. Transparent dental adhesive resin system (C&B SuperBond, Sun Medical).
11. Silicone impression material.
12. High-viscosity silicone oil.
13. Cotton swabs, tissue wipes, toothpicks, adhesive tape.

### 2.3 Surgical Instruments and General Equipment

1. Stereotaxic frame equipped with ear bars and bite bar.
2. Body temperature control system set at 37 °C.
3. Fiber light guide illumination.
4. Precision balance.
5. Iris scissors.
6. Fine forceps.
7. Scalpel handle with sterile blades.
8. Disposable syringes and needles.

### 2.4 Headplate and Fixation System

1. Custom stainless-steel headplate with a bilateral imaging chamber (Fig. 1g).
2. Custom head-fixation clamp compatible with the headplate (Fig. 1h).
3. Hex wrench or compatible fixation tool.

### 2.5 Macroscope Optical System

1. Two fast photographic lenses for tandem-lens macroscope configuration (e.g., 50 mm f/1.2 lenses).
2. Optional teleconverter for magnification adjustment.
3. Emission filter centered near 525 nm.
4. Dichroic mirror with cutoff near 490 nm.
5. 60-mm cage cube optical housing.
6. Adjustable optical support frame.

The overall optical layout is shown in Fig. 2a.

**Fig. 1.**
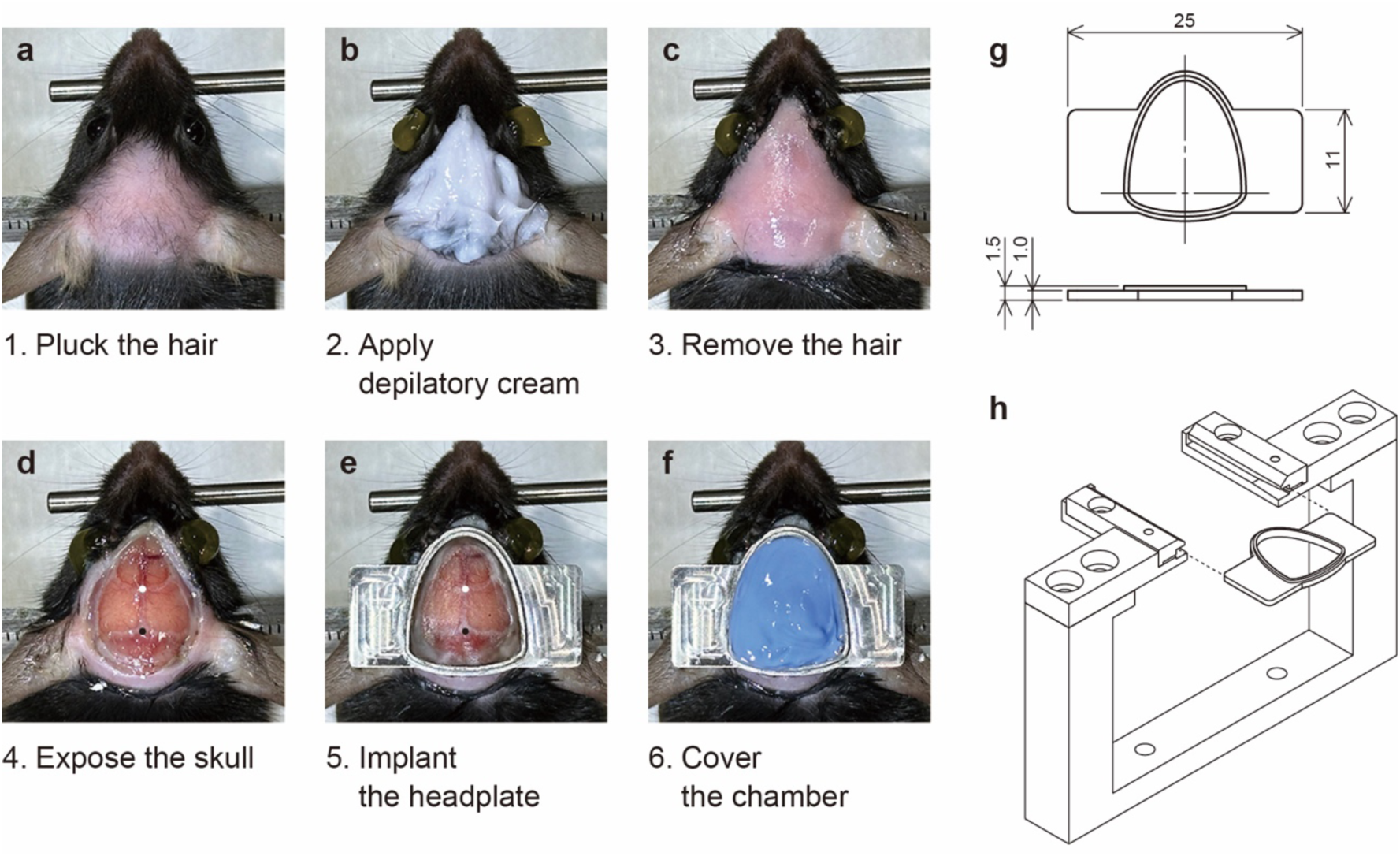
Surgical preparation, headplate design, and head-fixation system for chronic wide-field imaging in mice. (a) Hair on the scalp is manually plucked before chemical depilation. (b) Depilatory cream is applied to remove residual hair. (c) The scalp surface after complete hair removal. (d) The scalp is incised and retracted to expose the dorsal skull. White and black dots indicate reference points for stereotaxic alignment, bregma and lambda, respectively. (e) A custom stainless-steel headplate is positioned and fixed over the cleared skull. (f) The imaging chamber is temporarily filled with silicone impression material for postoperative protection or recovery. (g) Dimensioned drawing of the custom headplate (units: mm). (h) Assembled view of the custom head-fixation clamp used for stable imaging in living mice.

**Fig. 2.**
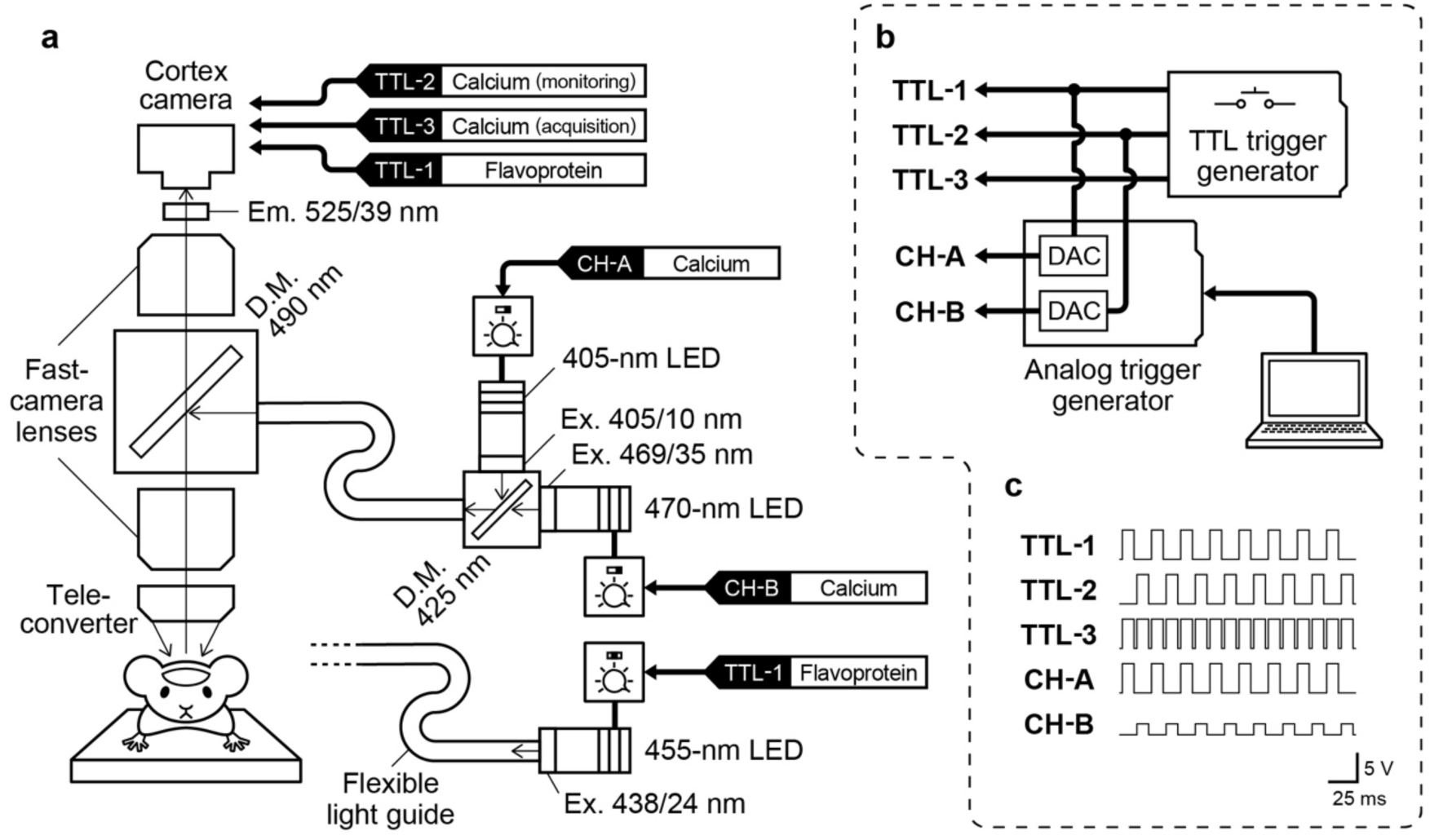
Optical configuration and trigger-control system for wide-field calcium and flavoprotein imaging. (a) Schematic diagram of the tandem-lens macroscope, illumination paths, and camera connections. Fluorescence emitted from the cortex passes through the emission path of a 490-nm dichroic mirror, is filtered through a 525/39-nm emission filter, and is recorded by the imaging camera. For calcium imaging, 405-nm and 470-nm LEDs are alternately passed through excitation filters, combined by a 425-nm dichroic mirror, and directed into a shared flexible light guide. For flavoprotein imaging, a 455- nm LED with an excitation filter is coupled to the same optical path. The excitation light is then reflected by a 490-nm dichroic mirror and delivered to the cortical surface through epi-illumination. Camera trigger inputs are switched according to imaging mode (calcium monitoring, calcium acquisition, or flavoprotein acquisition). (b) Block diagram of the custom trigger-control system. A transistor–transistor logic (TTL) trigger generator provides synchronized digital outputs (TTL-1 to TTL-3), while a computer-controlled analog trigger generator provides independently adjustable analog outputs (CH-A and CH-B) for LED intensity control and timing. DAC: digital-to-analog converter. (c) Representative timing waveforms for digital trigger signals and analog output channels. TTL-1, TTL-2, and TTL-3 control camera acquisition and synchronization, whereas CH-A and CH-B drive excitation LEDs for interleaved calcium imaging.

### 2.6 Illumination System

1. 405-nm LED excitation source.
2. 455-nm LED excitation source.
3. 470-nm LED excitation source.
4. LED drivers with external trigger input.
5. Excitation filter for 405-nm illumination.
6. Excitation filter for 435–455 nm illumination.
7. Excitation filter for 470-nm illumination.
8. Dichroic mirror with cutoff near 425 nm.
9. Collimating lens.
10. Achromatic relay lens.
11. Flexible light guide.
12. Optical table and dark enclosure.
13. Custom transistor–transistor logic (TTL) trigger generator.
14. Custom analog trigger generator with variable output amplitude. The typical trigger setting is shown in Figs. 2b, c.

### 2.7 Camera System

1. USB3 monochrome CMOS cameras with hardware trigger capability.
2. USB3 cables.
3. Trigger cables.
4. Infrared LED illumination for behavioral monitoring.
5. Computer workstation with sufficient RAM and SSD storage. Typical acquisition settings are summarized in Table 1.

### 2.8 Sensory Stimulation Devices

#### Visual Stimulation

1. Blue LED stimulus source.
2. Custom LED driver with digitally adjustable intensity.
3. Mechanical arm or positioning holder.

#### Somatosensory Stimulation

1. Miniature vibration motor for hindlimb stimulation.
2. Custom motor driver controlled by pulse-width modulation (PWM).

#### Control System

1. Microcontroller (Arduino)-based master controller.
2. MATLAB-based task control script for trial sequencing and synchronization. The overall mechanical layout and stimulus combinations are shown in Fig. 3.

### 2.9 Acquisition and Analysis Software

1. IC Express (The Imaging Source) or equivalent acquisition software capable of RAM-buffered recording and hardware-triggered acquisition.
2. MATLAB (MathWorks) for preprocessing, ΔF/F calculation, artifact reduction, trial extraction, block averaging, atlas registration, region-of-interest (ROI) analysis, and figure generation.
3. ImageJ/Fiji (optional) for image inspection and figure preparation.

Custom MATLAB scripts used in this chapter are available at: https://github.com/takamasa-yoshida/WFI-pipeline. The version associated with this chapter is archived at Zenodo: https://doi.org/10.5281/zenodo.20179799.

**Fig. 3.**
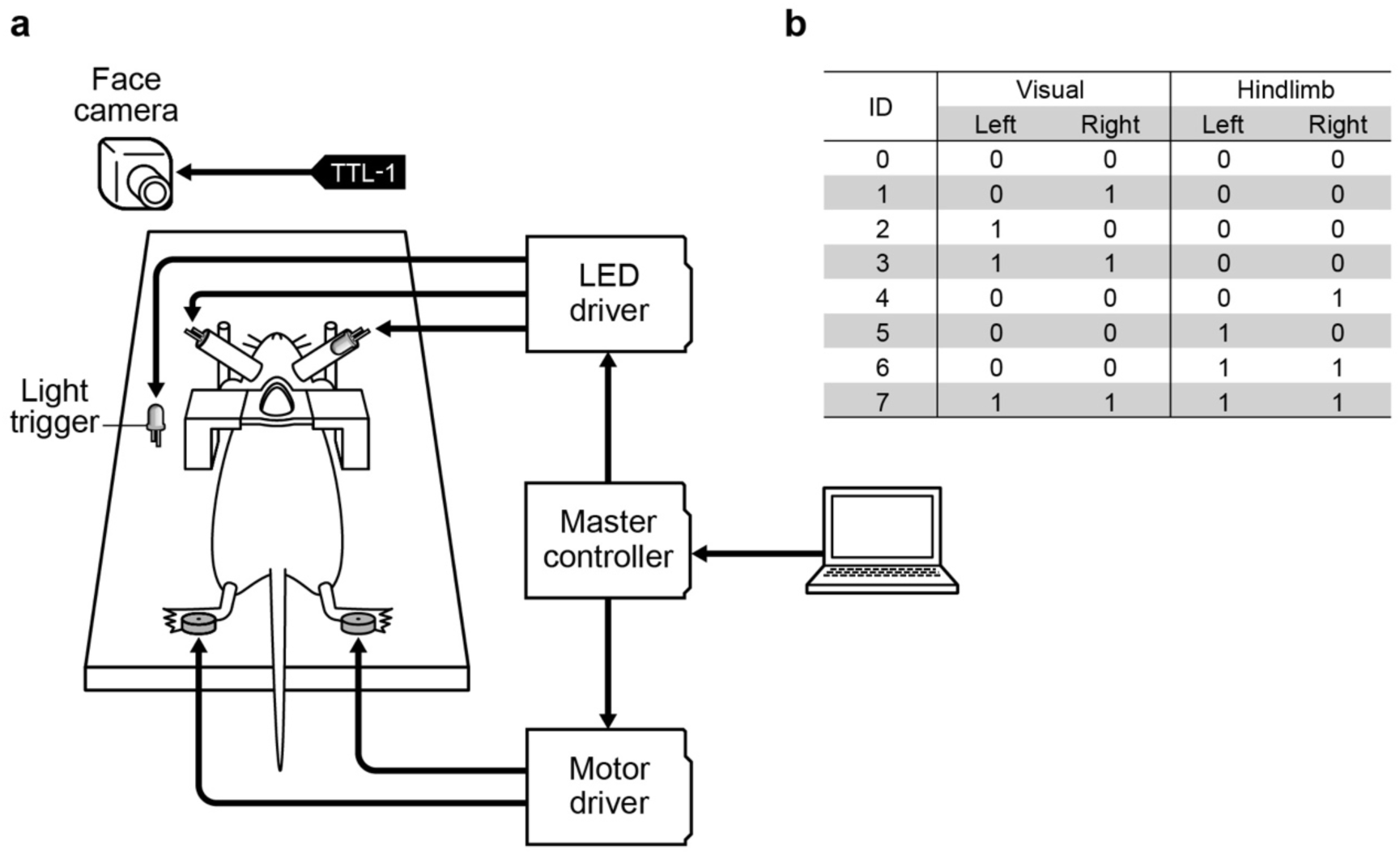
Sensory stimulation system and stimulus conditions used for multisensory wide-field imaging. (a) Schematic diagram of the stimulation setup. A custom master controller communicates with a computer and coordinates visual and somatosensory stimulation. Bilateral visual stimuli are delivered by LEDs positioned in close contact with the eyes through an LED driver. Somatosensory stimuli are delivered by miniature vibration motors attached to the hindlimbs through a motor driver. A light trigger LED placed near the animal provides a visual synchronization signal for the face camera. The face camera receives a synchronized TTL trigger signal for behavioral monitoring. (b) Stimulus combinations used in the experiment. Eight conditions were presented in pseudorandom order: no stimulus (ID 0), unilateral visual stimulation (IDs 1 and 2), bilateral visual stimulation (ID 3), unilateral hindlimb stimulation (IDs 4 and 5), bilateral hindlimb stimulation (ID 6), and combined bilateral visual plus bilateral hindlimb stimulation (ID 7).

**Table 1.** Recommended camera acquisition settings.

| Parameter | Calcium imaging | Flavoprotein imaging | Face camera |
| --- | --- | --- | --- |
| Camera | DMK33UX174 |  | DMK33UX273 |
| Frame rate | 80 frames/s<br>(interleaved mode) | 40 frames/s<br>(single mode) | 40 frames/s |
| Resolution | 1200 × 1200 |  | 640 × 480 (Binning 2x) |
| Exposure | 1/96 s | 1/47 s | 1/55 s |
| Gain | 24.0 dB | 25.4 dB | 0 dB |
| Trigger mode | External (trigger width) |  |  |
| Format | Y800 (8-bit grayscale) |  |  |

**Note:** Gain values should be adjusted depending on the imaging conditions.

## 3. Methods

### 3.1 Surgical Preparation

1. Weigh the mouse.
2. Prepare a mixed anesthetic solution to achieve final doses of 0.75 mg/kg medetomidine, 4 mg/kg midazolam, and 5 mg/kg butorphanol.
3. Administer the anesthetic intraperitoneally (see Notes 1 and 2).
4. Pluck the hair from the scalp using forceps (Fig. 1a; see Note 3).
5. Fix the mouse in the stereotaxic frame using the bite bar and ear bars (see Notes 4 and 5).
6. Adjust the bite bar vertically until the line connecting bregma and lambda is level.
7. Maintain body temperature at 37 °C throughout the procedure (see Note 6).
8. Apply ophthalmic ointment to both eyes (see Note 7).
9. Apply depilatory cream to the scalp for 2–3 min (Fig. 1b).
10. Remove the cream completely (Fig. 1c; see Note 8).
11. Disinfect the scalp and surrounding skin.
12. Apply lidocaine jelly to the incision site.
13. Incise and remove the scalp to expose the dorsal skull from the frontal region to the lambda suture (Fig. 1d).
14. Carefully remove periosteum and connective tissue from the skull surface (see Note 9).
15. Rinse the skull with sterile saline.
16. Dry the skull surface.

### 3.2 Skull Clearing and Headplate Implantation

1. Apply primer from the dental adhesive system to the exposed skull.
2. Incubate for 30–60 s.
3. Rinse thoroughly.
4. Prepare the resin monomer and catalyst according to the manufacturer’s instructions.
5. Apply the resin liquid across the skull surface (see Note 10).
6. Gradually build a transparent coating using alternating liquid and polymer applications.
7. Position the headplate parallel to the bregma–lambda axis with the chamber centered on the midline (Fig. 1e; see Note 11).
8. Seal all gaps between the skull and headplate using additional resin (see Note 12).
9. Allow the resin to fully cure.
10. Fill the chamber temporarily with silicone impression material if postoperative recovery is required before imaging (Fig. 1f; see Note 13).
11. Release the mouse from the stereotaxic frame.
12. Inject atipamezole hydrochloride to reverse medetomidine sedation if no consecutive imaging is planned.
13. Return the animal to a recovery cage until fully awake (see Note 14).

### 3.3 Imaging Setup

1. Secure the mouse attached to the headplate to the custom head-fixation clamp (Fig. 1h).
2. Remove the temporary chamber cover if used.
3. Fill the imaging chamber with high-viscosity silicone oil (see Notes 15 and 16).
4. Connect all trigger cables between LEDs, cameras, and trigger generators (Fig. 2a).
5. For calcium imaging, assign the 405-nm and 470-nm LEDs to alternating trigger channels and set the cortex camera to 80 frames/s for interleaved acquisition (Figs. 2b, c).
6. For flavoprotein imaging, connect the 455-nm LED and the cortex camera to the same trigger source and acquire at 40 frames/s (Figs. 2b, c).
7. Set the face camera to 40 frames/s (Fig. 3a).
8. Set LED drivers to external modulation or trigger mode.
9. Adjust LED output intensity as required (see Note 17).
10. Open the live camera preview.
11. Switch the trigger controller to monitoring mode for focusing.
12. For calcium imaging, temporarily set the cortex camera trigger to single-frame mode (40 frames/s) and synchronize acquisition with the 470 nm LED.
13. Adjust the Z-position of the macroscope until cortical blood vessels are sharply focused (Fig. 4a; see Note 18).
14. Confirm that no bubbles remain in the silicone oil layer.
15. Return the trigger configuration to acquisition mode before starting the experiment (see Notes 19 and 20).

### 3.4 Sensory Stimulation

1. Position the visual stimulus LEDs in close contact with the eyes using a shielded holder (Fig. 3a; see Note 21).
2. Attach vibration motors to the plantar surface of both hindlimbs using tape (Fig. 3a; see Note 22).
3. Place an optical trigger LED within the field of view of the face camera (Fig. 3a; see Note 23).
4. Program the stimulation controller to deliver eight multimodal stimulus conditions (Fig. 3b; see Note 24).
5. For calcium imaging, use a trial structure of 5 s prestimulus, 0.5 s stimulus, and 7 s poststimulus (12.5 s total; see Note 25).
6. Present all eight conditions in pseudorandom order for five blocks per set and repeat four sets (see Note 26).
7. For flavoprotein imaging, use a trial structure of 5 s prestimulus, 0.5 s stimulus, and 14.5 s poststimulus (20 s total; see Note 27).
8. Present all eight conditions in pseudorandom order for five blocks per set and repeat eight sets (see Note 28).

### 3.5 Data Acquisition

1. Launch the acquisition software.
2. Load the predefined camera settings (Table 1).
3. Specify a unique folder or prefix name for the recording set (see Note 29).
4. Prepare image capture in triggered mode.
5. Launch the MATLAB sequence-control interface.
6. Enter experimental parameters including animal ID, block number, and trial durations.
7. Start triggered image acquisition.
8. Immediately start the stimulation sequence.
9. Allow recording to continue until the programmed capture duration has ended.
10. Save all image sequences from RAM to disk immediately after frame acquisition (see Note 30).
11. Repeat additional sets as required using a new subfolder name for each set (see Note 31).
12. Verify that cortex images, face images, and stimulus log files were saved successfully (see Note 32).

### 3.6 Data Analysis

1. Import image sequences and stimulus log files into MATLAB.
2. Separate raw image streams into excitation-specific channels when alternating or interleaved acquisition was used.
3. Reconstruct temporally ordered image stacks for each channel (see Note 33).
4. Estimate slow baseline fluctuations for each pixel using a moving-window method.
5. Normalize image sequences to obtain relative signal changes (ΔF/F).
6. For calcium imaging datasets, regress the 405-nm reference signal from the 470-nm signal on a pixel-wise basis (see Note 34).
7. For flavoprotein datasets, perform baseline normalization and prestimulus correction.
8. Perform principal component analysis (PCA) when necessary to reduce the dimensionality of ΔF/F signals.
9. Apply independent component analysis (ICA) to separate putative neuronal and non-neuronal signal sources (see Note 35).
10. Remove selected non-neuronal independent components using a pixel-wise general linear model (GLM), using PCA-based regression as a fallback if ICA does not converge (see Note 36).
11. Segment data into peri-stimulus trials aligned to stimulus onset.
12. Average trials for each stimulus condition (Fig. 3b).
13. Generate stimulus-evoked activity maps from selected time windows (Fig. 4; see Note 37).
14. Register averaged maps or anatomical reference images to a standard cortical atlas (Fig. 4a; see Note 38).
15. Extract ROI-based temporal responses from predefined cortical regions (Figs. 4b–e).
16. Interpret delayed flavoprotein components cautiously (see Notes 39 and 40).

The computational workflow has been described in related wide-field imaging protocols [4,5].

**Fig. 4.**
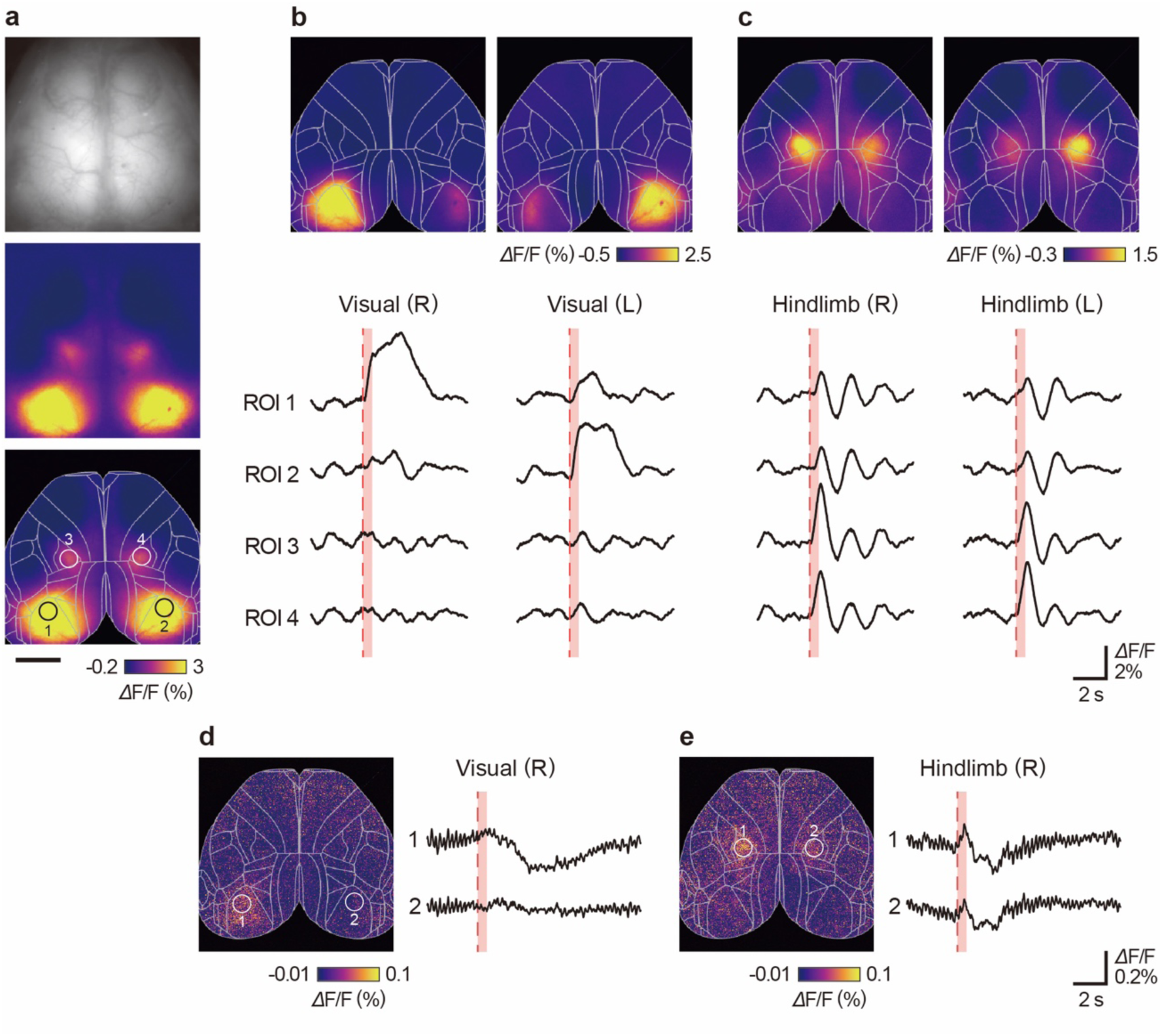
Representative stimulus-evoked responses and ROI-based signal analysis obtained by wide-field calcium and flavoprotein imaging. (a) Example calcium-imaging dataset during combined bilateral visual plus bilateral hindlimb stimulation (stimulus ID 7). Raw cortical image (top), stimulus-evoked activity map (middle), and atlas- registered map with representative regions of interest (ROIs; bottom). Registration was performed by matching the four peak response locations to bilateral primary visual cortex (V1) and hindlimb primary somatosensory cortex (S1HL) in the Allen Brain Atlas using scaling, rotation, and translation. ROIs 1 and 2 indicate bilateral V1, and ROIs 3 and 4 indicate bilateral S1HL. Scale bar, 2 mm. (b, c) Calcium responses to unilateral visual (b) and hindlimb (c) stimulation. Top panels show activity maps for right and left stimulation, calculated as the mean response from stimulus offset to 1 s post- offset. Lower traces show averaged ΔF/F signals from ROIs 1–4. Contralateral-dominant responses are observed in both conditions. (d, e) Flavoprotein autofluorescence responses to right visual (d) and right hindlimb (e) stimulation. Each panel shows the activity map with corresponding bilateral ROIs and averaged ΔF/F traces. Activity maps were calculated as the mean response from 125 ms after stimulus onset over a 500-ms window. Lower traces show averaged ΔF/F signals from ROIs 1 and 2. Delayed flavoprotein signals may include vascular and glial metabolic components and should be interpreted cautiously. Red shaded areas in the traces indicate stimulus presentation periods in panels b–e.

## 4. Notes

1. Adjust anesthetic volume according to body weight immediately before injection.
2. Confirm adequate anesthesia by loss of pedal withdrawal reflex (toe pinch) before surgery.
3. Perform rough hair removal before ear-bar fixation to reduce accidental injury during handling.
4. The distance between ear bars should not be smaller than the interaural distance (typically 10– 11 mm in adult mice).
5. Excessive ear-bar pressure may damage the skull base or cause pain responses.
6. Maintain body temperature throughout surgery and imaging sessions whenever anesthesia is used.
7. Applying ophthalmic ointment prevents corneal drying.
8. Remove all depilatory cream with a water-moistened cotton swab to prevent skin irritation.
9. Ensure complete removal of residual tissue by gentle scraping with a scalpel before resin coating to improve adhesion and optical clarity.
10. Applying the monomer solution first improves the adhesion of the polymer to the skull surface, resulting in increased transparency and mechanical stability.
11. The headplate should be placed to maximize bilateral cortical coverage.
12. Incomplete sealing around the headplate may cause leakage of silicone oil during imaging.
13. Silicone impression material helps protect the transparent skull surface from dust and mechanical damage.
14. Minor intracranial hemorrhage observed immediately after surgery may resolve spontaneously after several days of recovery.
15. Remove air bubbles from the silicone oil layer using a toothpick or by allowing the preparation to stand for approximately 20 min.
16. Prevent hair around the headplate from contacting the silicone oil surface inside the chamber, as this may cause leakage due to capillary wicking.
17. If illumination power is insufficient, use a dual-branch bundle fiber light guide to combine output from two independent light-source systems.
18. A teleconverter mounted in the reverse orientation can be used to obtain lower magnification and a wider field of view (e.g., a 1.4× teleconverter provides approximately 0.71× magnification).
19. For calcium imaging, verify that 405-nm and 470-nm frames are correctly interleaved and temporally aligned before starting formal acquisition.
20. If triggering fails, first confirm trigger mode settings and then verify exposure time constraints relative to frame rate (Table 1). The exposure time must not exceed the reciprocal of the frame rate.
21. The visual stimulus LEDs had a center wavelength of 468 nm and an irradiance of approximately 3.0 mW/m², delivered as a 10-Hz square-wave flicker. Blue light was selected to avoid overlap with the fluorescence emission band while still evoking robust responses in mouse visual cortex [38].
22. Hindlimb stimulation was delivered using a disk-shaped vibration motor (10 mm diameter, 3.3 mm thickness; LBV10B-009, Nidec), producing approximately 170 Hz sinusoidal vibration.
23. The light trigger signal was presented simultaneously during stimulus onset to synchronize stimulus timing with cortex image acquisition.
24. Verify that all stimulus devices are functioning normally before recording.
25. For calcium imaging, avoid trial repetition rates close to strong vasomotion frequencies (∼0.1 Hz).
26. The number of sets depends on the available RAM capacity of the acquisition computer.
27. For flavoprotein imaging, use sufficiently long intertrial intervals to allow complete recovery of metabolic signals.
28. Flavoprotein imaging generally requires a higher number of trial repetitions than calcium imaging to obtain a sufficient signal-to-noise ratio.
29. Reassign a unique recording prefix or folder name for each imaging set.
30. Use SSD storage for high-speed acquisition and maintain sufficient free space (>50%) to avoid decreases in data transfer speed.
31. Set the recording duration to fully utilize available RAM and acquire data in multiple runs to reduce the risk of data loss.
32. Save raw image files together with stimulus log files and behavioral recordings into a common folder to facilitate later reanalysis.
33. For interleaved acquisition, temporal offsets between alternating frames should be corrected by applying pixel-wise linear interpolation.
34. Pixel-wise regression of the 405-nm reference channel can improve removal of shared hemodynamic and motion-related artifacts.
35. Select non-neuronal components based on their spatial and temporal characteristics, such as signals outside the cortical mask, spatially uniform patterns, or slow, stimulus-independent fluctuations. ICA may be particularly useful for flavoprotein imaging, where vascular and global components are prominent.
36. Use the selected component time series as nuisance regressors in a pixel-wise GLM and subtract their contributions from the original ΔF/F signals. If ICA does not converge, reduce the number of components and, if necessary, use selected PCA components directly for GLM-based correction.
37. The time window used for stimulus-evoked maps should be selected according to signal kinetics.
38. When registering activity maps to the Allen Brain Atlas [39], use robust anatomical landmarks (e.g., skull suture patterns) and reproducible sensory-response peaks (e.g., bilateral V1 and S1HL) for alignment.
39. The delayed component of flavoprotein autofluorescence should be interpreted cautiously, as it may reflect not only hemodynamic effects but also slower neurometabolic processes involving glial as well as neuronal compartments [15].
40. For flavoprotein imaging, separation of hemodynamic components should be considered using multi-wavelength simultaneous measurements [40,41].

## Acknowledgement

This work was supported by JSPS KAKENHI Grant Number JP23K06004.

